# Rampant C->U hypermutation in the genomes of SARS-CoV-2 and other coronaviruses – causes and consequences for their short and long evolutionary trajectories

**DOI:** 10.1101/2020.05.01.072330

**Authors:** P. Simmonds

**Affiliations:** Nuffield Department of Medicine, University of Oxford, South Parks Road, Oxford, OX1 3SY, UK

## Abstract

The pandemic of SARS coronavirus 2 (SARS-CoV-2) has motivated an intensive analysis of its molecular epidemiology following its worldwide spread. To understand the early evolutionary events following its emergence, a dataset of 985 complete SARS-CoV-2 sequences was assembled. Variants showed a mean 5.5-9.5 nucleotide differences from each other, commensurate with a mid-range coronavirus substitution rate of 3×10^−4^ substitutions/site/year. Almost half of sequence changes were C->U transitions with an 8-fold base frequency normalised directional asymmetry between C->U and U->C substitutions. Elevated ratios were observed in other recently emerged coronaviruses (SARS-CoV and MERS-CoV) and to a decreasing degree in other human coronaviruses (HCoV-NL63, -OC43, -229E and -HKU1) proportionate to their increasing divergence. C->U transitions underpinned almost half of the amino acid differences between SARS-CoV-2 variants, and occurred preferentially in both 5’U/A and 3’U/A flanking sequence contexts comparable to favoured motifs of human APOBEC3 proteins. Marked base asymmetries observed in non-pandemic human coronaviruses (U>>A>G>>C) and low G+C contents may represent long term effects of prolonged C->U hypermutation in their hosts.

**Importance:** The evidence that much of sequence change in SARS-CoV-2 and other coronaviruses may be driven by a host APOBEC-like editing process has profound implications for understanding their short and long term evolution. Repeated cycles of mutation and reversion in favoured mutational hotspots and the widespread occurrence of amino acid changes with no adaptive value for the virus represents a quite different paradigm of virus sequence change from neutral and Darwinian evolutionary frameworks that are typically used in molecular epidemiology investigations.

## INTRODUCTION

SARS-coronavirus-2 (SARS-CoV-2) emerged late 2019 in the Hubei province, China as a cause of respiratory disease occasionally leading to acute respiratory distress syndrome and death (COVID-19)(1-4). Since the first reports in December, 2019, infections with SARS-CoV-2 have been reported from a rapidly increasing number of countries worldwide, and led to its declaration as a pandemic by the World Health Organisation in March, 2020. In order to understand the origins and transmission dynamics of SARS-CoV-2, sequencing of SARS-CoV-2 directly from samples of infected individuals worldwide has been performed on an unprecedented scale. These efforts have generated many thousands of high quality consensus sequences spanning the length of the genome and have defined a series of geographically defined clusters that recapitulate the early routes of international spread. However, as commented elsewhere (https://doi.org/10.1101/2020.03.16.20034470), there is remarkable little virus diversity at this early stage of the pandemic and analyses of its evolutionary dynamics remain at an early stage.

The relative infrequency of substitutions is the consequence of a much lower error rate on genome copying by the viral RNA polymerase of the larger nidovirales, including coronaviruses. This is achieved through the development of a proofreading capability through mismatch detection and excision by a viral encoded exonuclease, Nsp14-ExoN (5-7). Consequently, coronaviruses show a low substitution rate over time, typically in the range of 1.5 – 10 × 10^−4^ substitutions per site per year (SSY) (8-13). Applying a mid-range estimate to the 3-5 month timescale of the SARS-CoV-2 pandemic indicates that epidemiologically unrelated strains might show around 6-10 nucleotides differences from each other over the 30,000 base length of their genomes.

In the current study, we have analysed the nature of the sequence diversity generated within the SARS-CoV-2 virus populations revealed by current and ongoing virus sequencing studies. We obtained evidence for a preponderance of driven mutational events within the short evolutionary period following the zoonotic transmission of SARS-CoV-2 into humans. Sequence substitutions were characterised by a preponderance of cytidine to uridine (C->U) transitions. The possibility that the initial diversity within a viral population was largely host-induced would have major implications for evolutionary reconstruction of SARS-CoV-2 variants in the current pandemic, as well as in our understanding both of host antiviral pathways against coronaviruses and the longer term shaping effects on their genome composition.

## RESULTS

### Sequence changes in SARS-CoV-2

Four separate datasets of full length, (near-) complete genome sequences of SARS-CoV-2 sequences collected from the start of the pandemic to those most recently deposited on the 24^th^ April, 2020 were aligned and analysed. Each dataset showed minimal levels of sequence divergence, with mean pairwise distances ranging from 5.5-9.5 nucleotide differences between each sequence. However, several aspects of the frequencies and sequence contexts of the observed changes were unexpected. Firstly, the ratio of non-synonymous (amino acid changing) to synonymous substitutions was high – in the range of 0.57-0.73 amongst the different SARS-CoV-2 datasets. This contrasts with a much lower ratio (consistently below <0.22) in sequence datasets assembled for the other human coronaviruses (Table 1). Including a range of coronaviruses in the analysis, there was a consistent association between dN/dS with degree of sequence divergence (Fig. 1).

**TABLE 1.** CORONAVIRUS SEQUENCE DATASETS USED FOR THE STUDY.

CORONAVIRUS SEQUENCE DATASETS USED FOR THE STUDY
| **Virus** | **Family** | **Number** | **Length** | **MPD<sup>1</sup>** | **dN/dS<sup>2</sup>** |
| --- | --- | --- | --- | --- | --- |
| *<u>Zoonotic coronaviruses</u>* |  |  |  |  |  |
| SARS-CoV-2_Charite | *Coronaviridae* | 115 | 29748 | 0.000187 | 0.728 |
| SARS-CoV-2_Repl1 | *Coronaviridae* | 300 | 29409 | 0.000267 | 0.569 |
| SARS-CoV-2_Repl2 | *Coronaviridae* | 300 | 29408 | 0.000306 | 0.630 |
| SARS-CoV-2_Repl3 | *Coronaviridae* | 286 | 29404 | 0.000322 | 0.650 |
| SARS-CoV-1-like (bat) | *Coronaviridae* | 40 | 29480 | 0.048414 | 0.121 |
| SARS-CoV-1 | *Coronaviridae* | 22 | 29443 | 0.000381 | 0.428 |
| MERS-CoV | *Coronaviridae* | 26 | 30043 | 0.005065 | 0.228 |
| *<u>Other human and related coronaviruses</u>* |  |  |  |  |  |
| OC43-human | *Coronaviridae* | 178 | 30135 | 0.0081 | 0.219 |
| OC43-bovine | *Coronaviridae* | 113 | 30485 | 0.0104 | 0.139 |
| HKU1-gt1 | *Coronaviridae* | 27 | 29613 | 0.0023 | 0.155 |
| HKU1-gt2 | *Coronaviridae* | 12 | 29610 | 0.0121 | 0.096 |
| NL63 | *Coronaviridae* | 61 | 27453 | 0.0086 | 0.131 |
| 229E-human | *Coronaviridae* | 26 | 26846 | 0.0071 | 0.203 |
| 229E-camel | *Coronaviridae* | 33 | 27051 | 0.002 | 0.167 |

<sup>1</sup>Mean nucleotide p distances between complete genome sequences
<sup>2</sup>Frequency of non-synonymous (dN) to synonymous (dS) p distances

**FIGURE 1.**
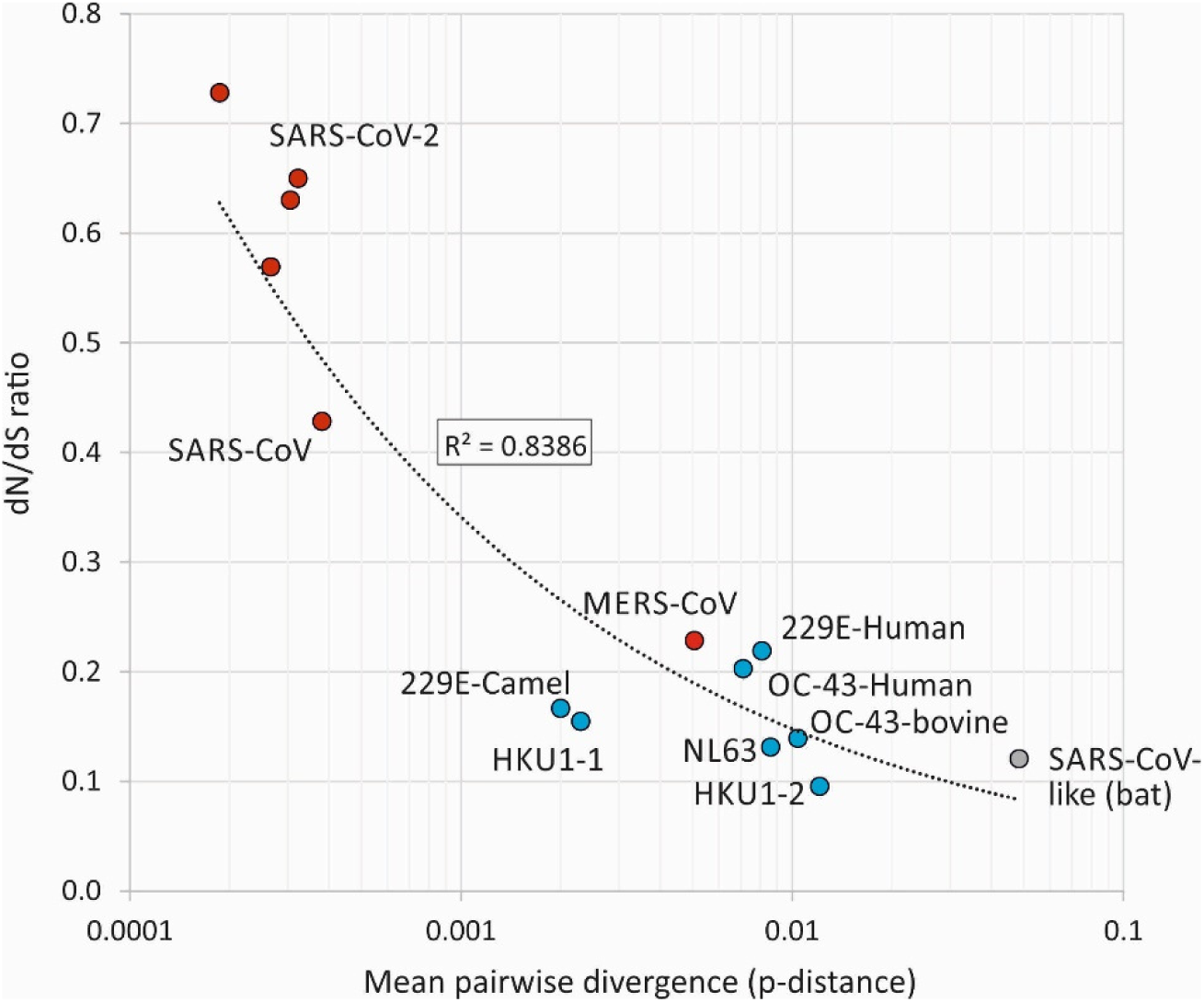
ASSOCIATION BETWEEN SEQUENCE DIVERGENCE AND dN/dS RATIO. A comparison of dN/dS ratios in recently emerged coronaviruses (red circles), other human coronaviruses and relatives infecting other species (blue circles) and a collection of bat SARS-like viruses (grey circle). A power law line of best fit showed a significant correlation between divergence and dN/dS ratio (*p* = 0.000006).

We next estimated frequencies of individual transitions and transversions occurring during the short-terms evolution of SARS-CoV-2. Sequence differences between each SARS-CoV-2 full genome sequence and a majority rule consensus sequence generated for each of the four SARS-CoV-2 datasets were calculated. The directionality of sequence change underlying the observed substitutions was inferred through restricting analysis to polymorphic sites with a minimal number of variable bases (typically singletons). In practice because of the scarcity of substitutions, variability thresholds of 10%, 5%, 2% and 1% yielded similar numbers and relative frequencies of each transition and transversion. Equivalent evidence for directionality was obtained through comparison of each sequence in the dataset with the first outbreak sequence (MN908947; Wuhan-Hu-1), approximately ancestral to current circulating SARS-CoV-2 strains (data not shown). For the purposes of the analysis presented here, a consensus-based 5% threshold was used.

A listing of the sequence changes revealed a striking (approximately four-fold) excess of sites where C->U substitutions occurred in SARS-CoV-2 sequences compared to the other three transitions (Fig. 2A). This excess was the more remarkable given there was an almost two-fold greater number of U bases in the SARS-CoV-2 genome compared to C’s (32.1% compared to 18.4%). To formally analyse the excess of C->U transitions we calculated an index of asymmetry (frequency[C->U] / f[U->C]) x (fU/fC) and compared this with degrees of sequence divergence and dN/dS ratio in SARS-CoV-2 and other coronavirus datasets (Fig. 2B, 2C). This comparison showed that the excess of C->U substitutions was most marked among very recently diverged sequences associated with the SARS-CoV-2 and SARS-CoV outbreaks and was reduced significantly in sequence datasets of the more divergent human coronaviruses (NL63, OC43, 229E and OC43) datasets as sequences accumulated substitutions.

**FIGURE 2.**
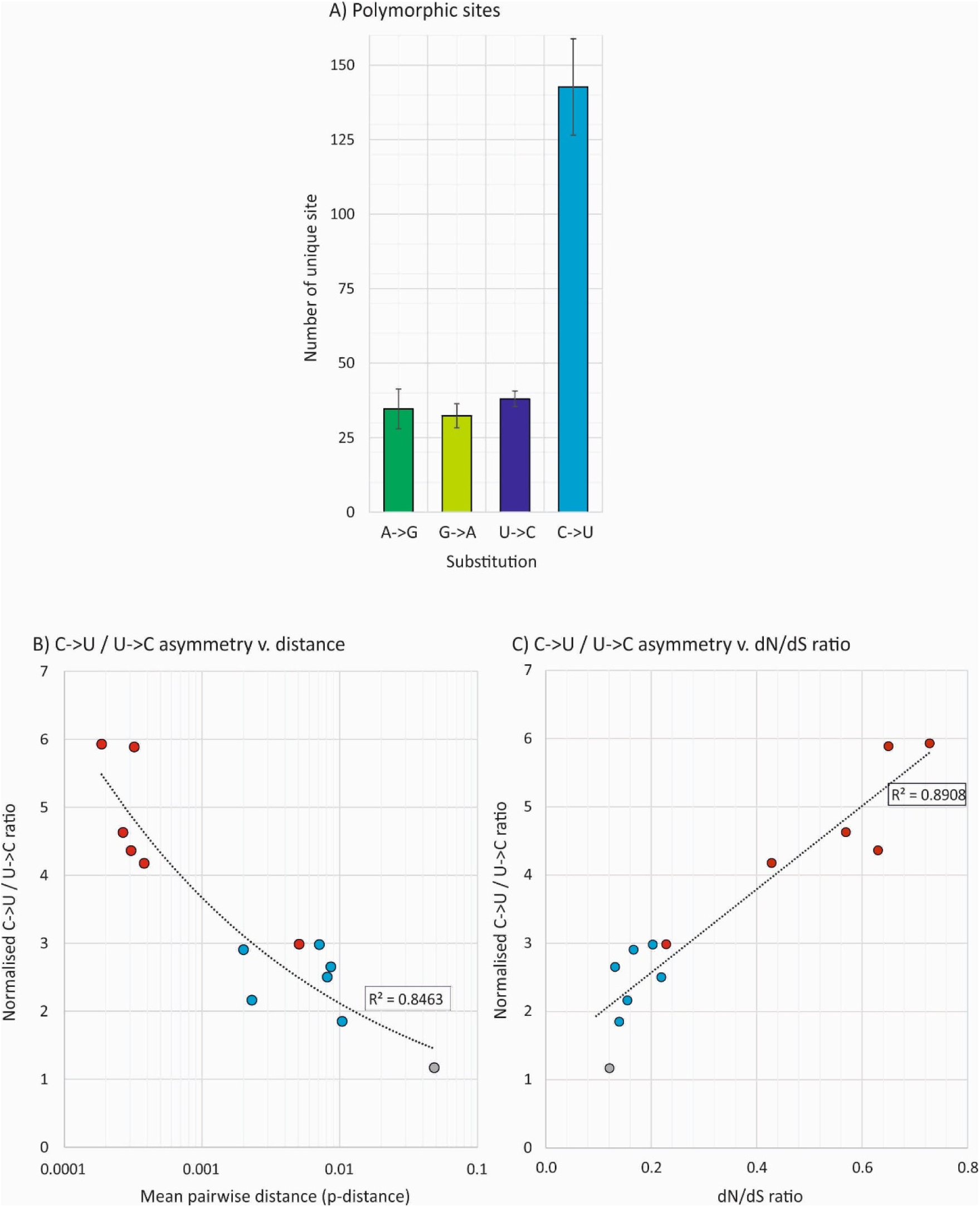
ASSOCIATION OF EXCESS C->U TRANSITIONS WITH DIVERGENCE. A) Numbers of sites in the SARS-CoV-2 genome with each of the four transitions. Bar heights represent the mean of the three sequence samples; error bars show one standard deviation (SD). (B) Relationship between sequence diversity and a normalised metric of asymmetry between the numbers of C->U and U > C transitions (where 1.0 is the expected number). Power law regression line was significant at *p* < 0.0001. (C) Association of dN/dS ratio with C->U / U > C asymmetry. The power law regression lines were significant at *p* = 0.001 and 0.0004 respectively. Points are coloured as in Fig. 1.

C->U substitutions were scattered throughout the SARS-CoV-2 genome (Fig. 3). Long bars representing more polymorphic sites were frequently shared between replicate datasets but unique substitutions (occurring once in the dataset - short bars) showed largely separate distributions. Substitutions were not focussed towards any particular gene or intergenic region although all three datasets showed marginally higher frequencies of substitutions in the N gene. The grouping of a selection of sequences showing C->U changes in different genome regions were plotted in a phylogenetic tree containing sequences from the SARS-CoV-2 dataset (Fig. 4). Within the resolution possible in tree generated from such a minimally divergent dataset, many sequences with shared C-U changes were not monophyletic (*eg*. those with substitutions at positions 5784, 10319, 21575, 28657 and 28887). This lack of grouping is consistent with multiple *de novo* occurrences of the same mutation in different SARS-CoV-2 lineages.

**FIGURE 3.**
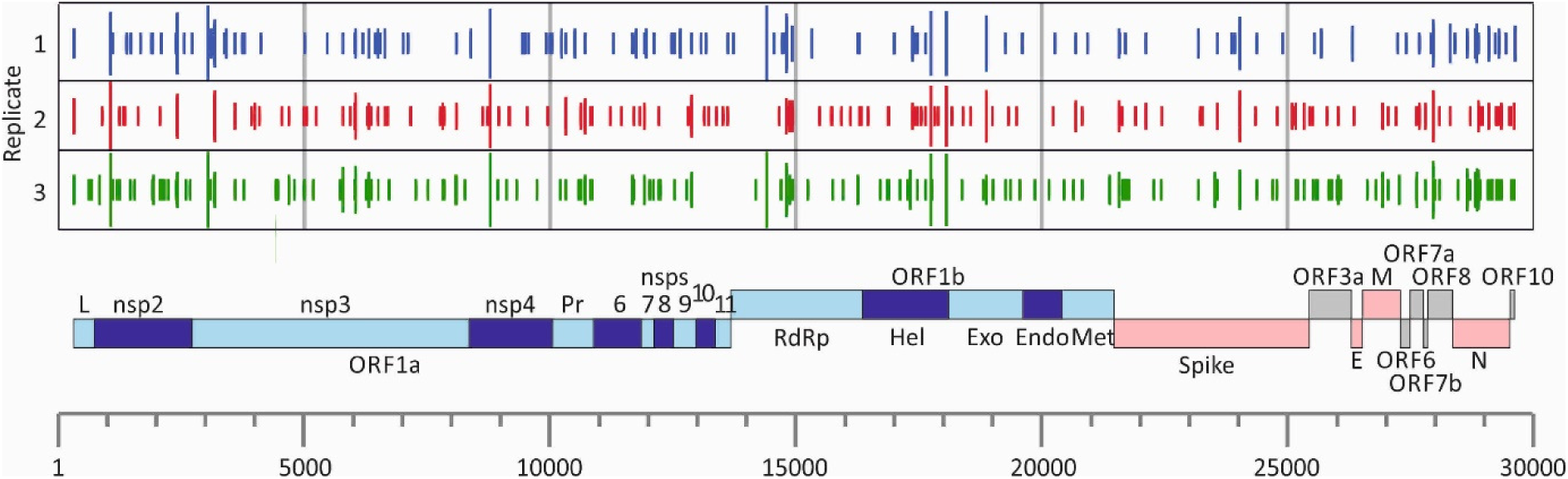
POSITIONS OF C->U TRANSITIONS IN THE SARS-CoV-2 GENOME. Positions of C->U substitutions in each of the three replicate SARS-CoV-2 sequence datasets, matched to a genome diagram of SARS-CoV-2 (using the annotation from the prototype sequence MN908947). The number of transitions at each site are shown on a log scale, with the shortest bars indicating individual substitutions.

**FIGURE 4.**
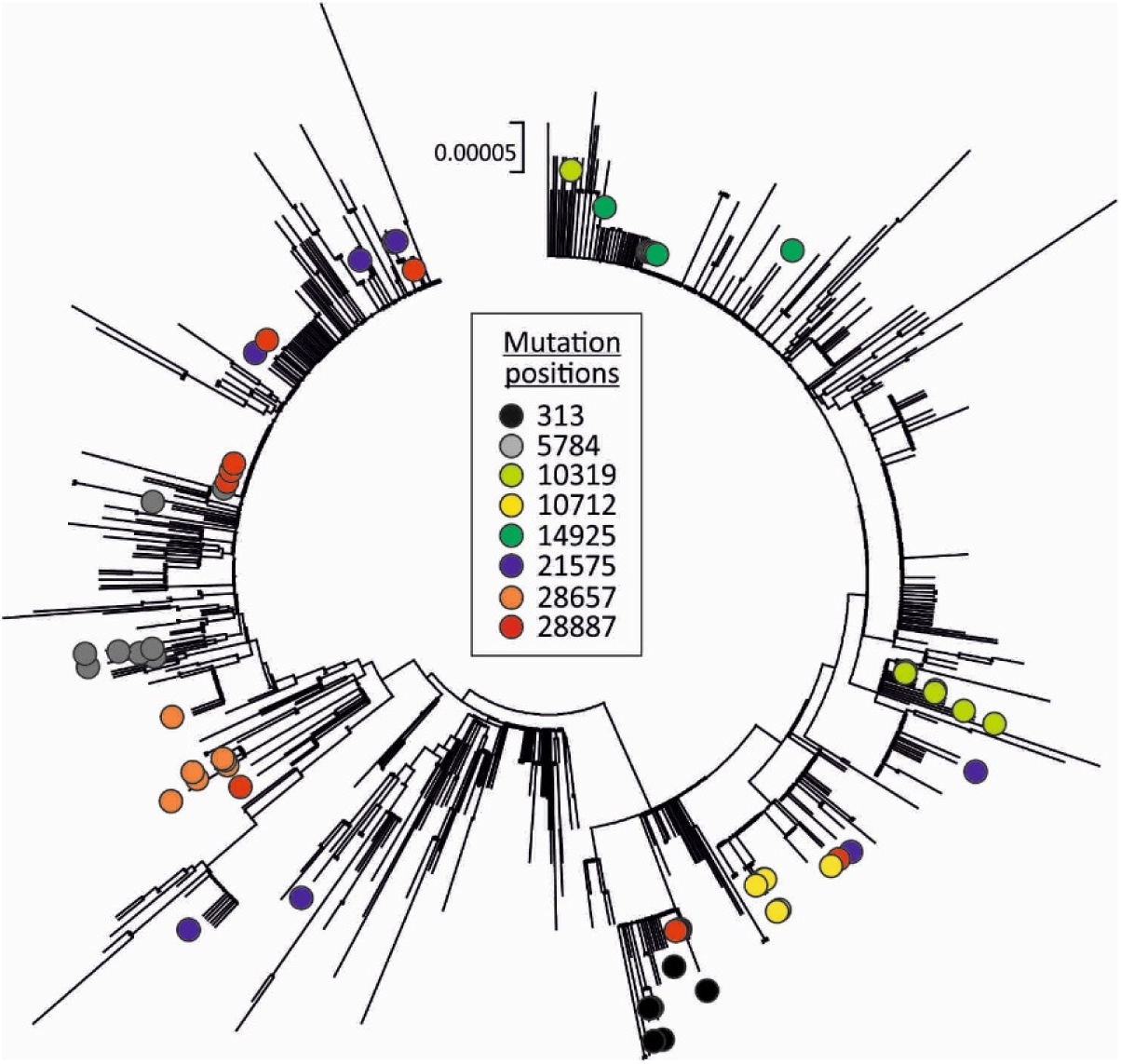
PHYLOGENY OF SARS-CoV-2 AND POSITIONS OF SEQUENCES WITH C->U CHANGES. Neighbour-joining tree of 865 SARS-CoV-2 complete genome sequences constructed in MEGA6 (37). Labels show the position of sequences containing a selection of C->U transitions at the genome positions indicated in the key.

The abnormally high dN/dS ratios of 0.6-0.7 in SARS-CoV-2 sequences (Table 1; Fig. 1) indicated that around 50% of nucleotide substitutions would produce amino acid changes (if approximately 75% of nucleotide changes are non-synonymous). On analysis of amino acid sequence changes, a remarkable 50% of non-synonymous substitutions in the SARS-CoV-2 sequence dataset were the consequence of C->U transitions (Fig. 5). The underlying mechanism that leads to C->U hypermutation therefore also drives much of the amino acid sequence diversity observed in SARS-CoV-2.

**FIGURE 5.**
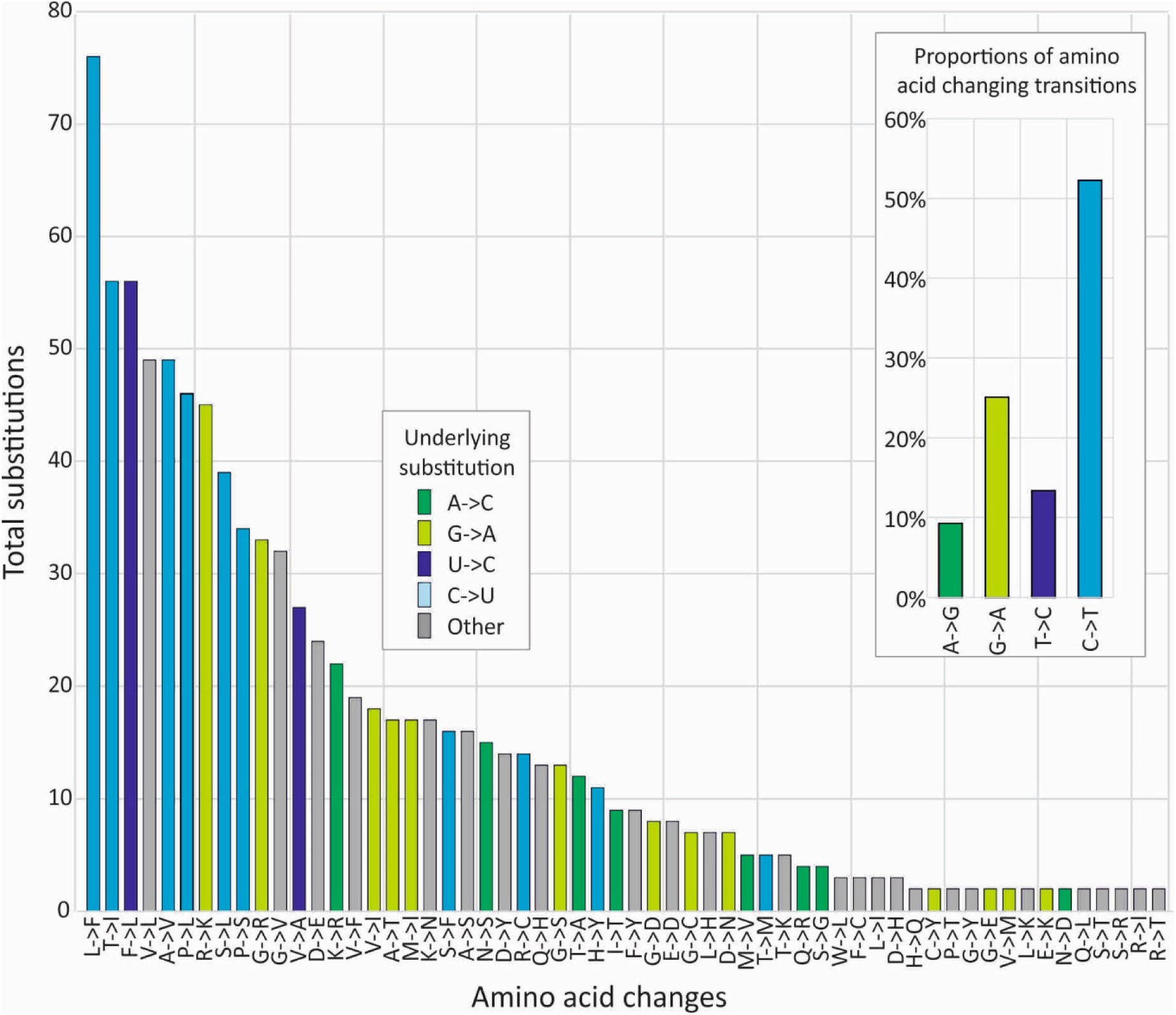
AMINO ACID CHANGES INDUCED BY DIFFERENT NUCLEOTIDE SUBSTITUTIONS. Numbers of individual amino acids changes observed in the combined SARS-CoV-2 dataset (864 sequences) at a 5% variability threshold. Bars are coloured based on the underlying nucleotide changes. Inset graph shows the relative proportions of transitions leading to amino acid changes.

The context of cytidines within a sequence strongly influenced the likelihood of it mutating to a U (Fig. 6). The greatest numbers of mutations were observed if the upstream (5’) base was an A or U. There was also a similar approximately 4-fold increase in transitions if these bases were located on the downstream (3’) side. The effects of the 5’ and 3’ contexts were additive; C residues surrounded by an A or U at both 5’ and 3’ sides were 10-fold more likely to mutate than those flanked by C or G residues (mean 31.9 transitions compared to 3.6). Splitting the data down into the 16 combinations of 5’ and 3’ contexts, a 5’U far more potently restricted non-C->U substitutions than a 5’A (Fig. S1; Suppl. Data), while 5’G or 5’C almost eliminated substitutions irrespective of the 3’ context. No context created any substantial asymmetry in G->A compared to A->G transitions.

**FIGURE 6.**
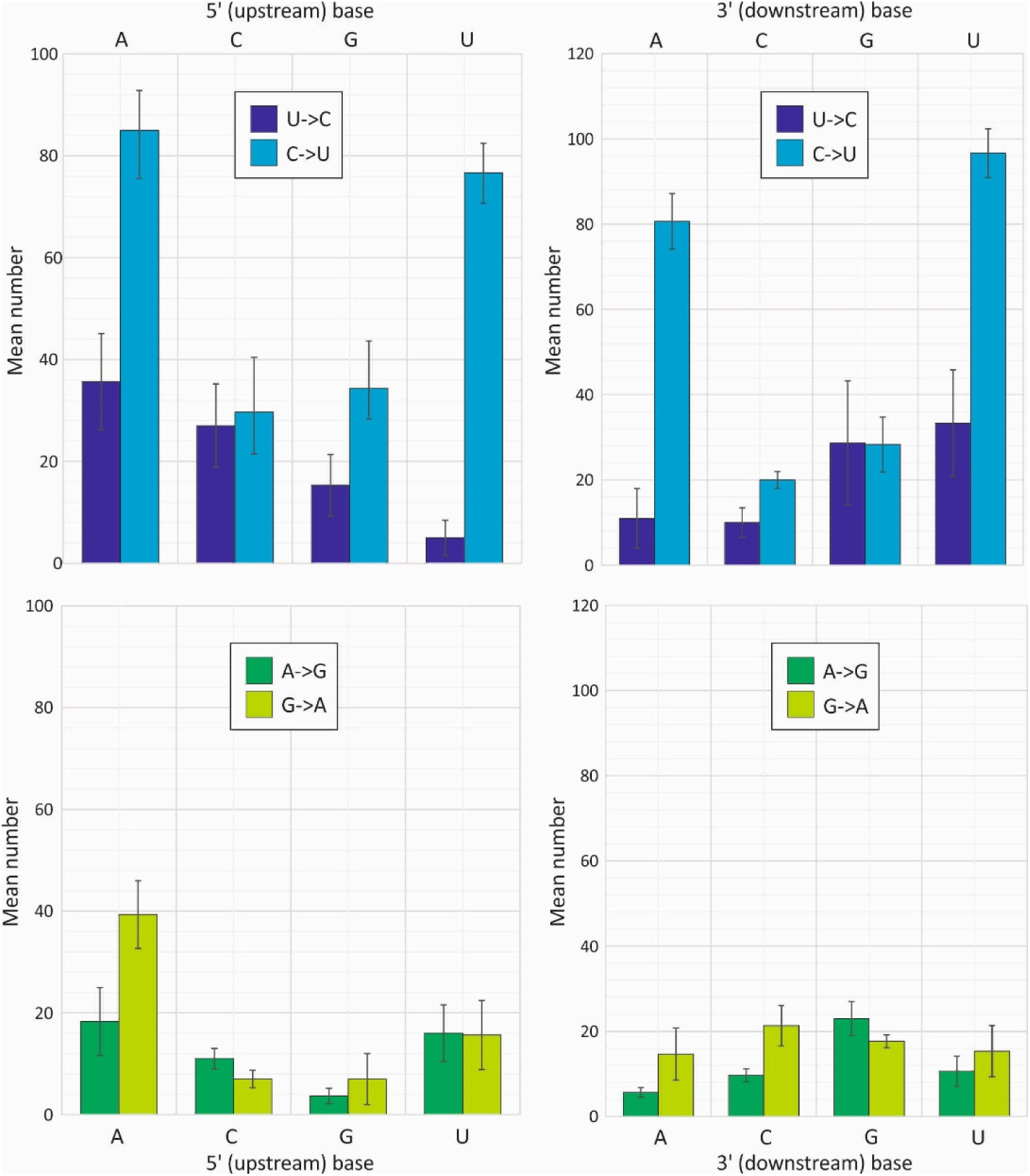
INFLUENCE OF 5’ AND 3’ BASE CONTEXTS ON C/U AND G/A TRANSITION FREQUENCIES. Totals of each transition in the SARS-CoV-2 sequence dataset split into sub-totals based on the identity of the 5’ (left) and 3’ (right) base. Bar heights represent the mean of the three sequence samples; error bars show standard deviations. A further division into the 16 combinations of 5’ and 3’ base contexts is provided in Fig. S1; Suppl., Data.

The G+C content of coronaviruses varied substantially between species, with highest frequencies in the recently emerged zoonotic coronaviruses (MERS-CoV: 41%, SARS-CoV: 41% and SARS-CoV-2: 38%) and lowest in HKU1 (32%). Collectively, there was a significant relationship between C depletion and U enrichment with G+C content (Fig. 7). The difference in G+C content was indeed almost entirely attributable to changes in the frequencies of C and U bases; the 9% difference in G+C content between MERS-CoV and HKU1 arose primarily from the 20% -> 13% reduction in frequencies of C. There was a comparable 8% increase in the frequency of U. Their combined effects left frequencies of G and A relatively unchanged. It appears that the accumulated effect of the C->U / U->C asymmetry led to marked compositional abnormalities in coronaviruses.

**FIGURE 7.**
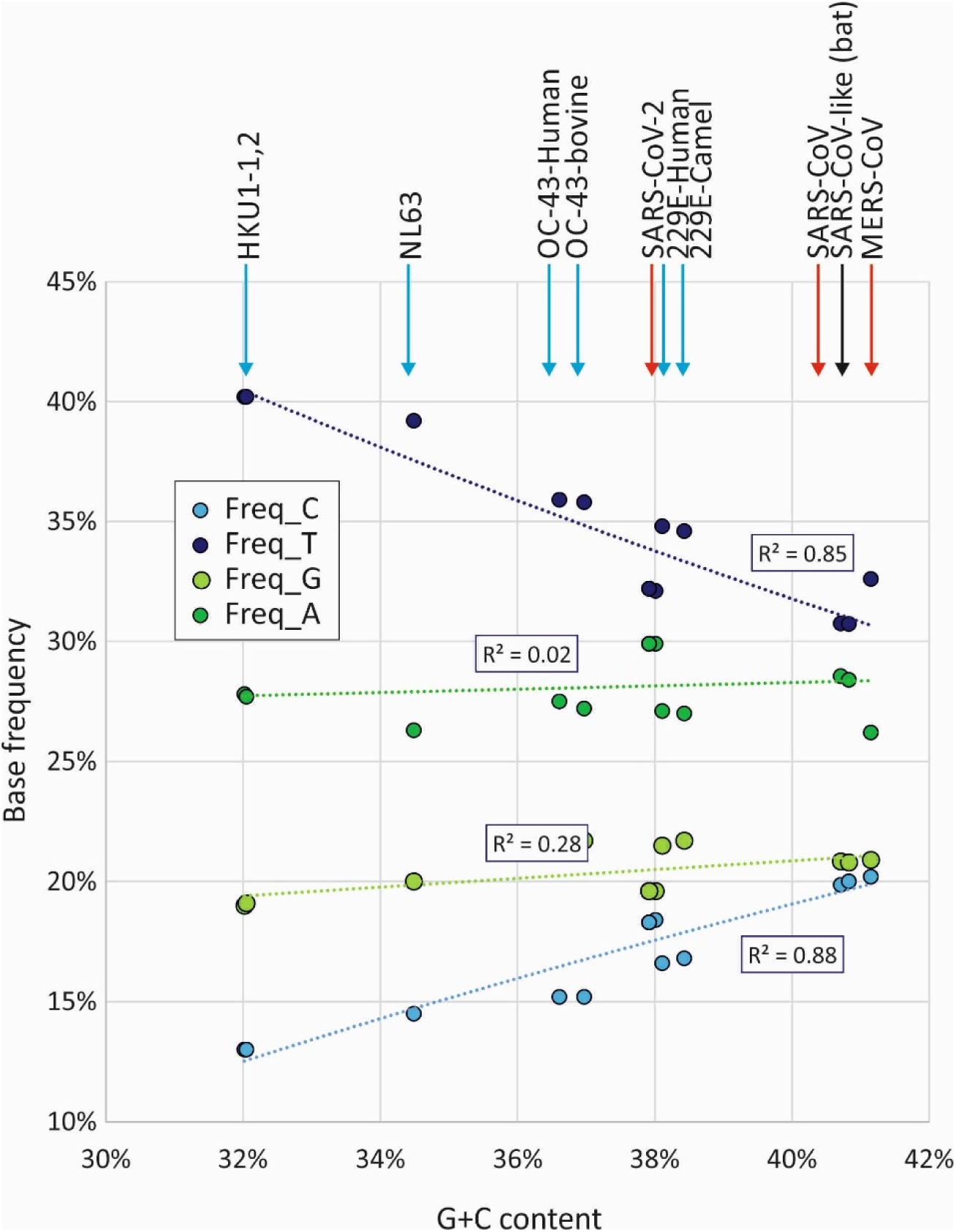
BASE FREQUENCIES IN DIFFERENT CORONAVIRUSES. Relationship between G+C content and frequencies of individual bases in coronaviruses. The associations between C depletion and U enrichment with G+C content were both significant by linear regression at *p* = 5 × 10^−7^ and *p* = 5 × 10^−6^ respectively. No significant associations were observed between G+C content and G (*p* = 0.05) or A (*p* = 0.62) frequencies. Arrows are colour coded as for Fig. 1.

## DISCUSSION

The wealth of sequence data generated since the outset of the SARS-CoV-2 pandemic, the accuracy of the sequences obtained by a range of NGS technologies and the long genomes and very low substitution rate of coronaviruses provided a unique opportunity to investigate sequence diversification at very high resolution. The findings additionally provide insights into the mutational mechanisms and contexts where sequence changes occur. Thirdly, it informs us about the longer term evolution of viruses and potential effects of the cell in moulding virus composition.

### The mechanism of SARS-CoV-2 hypermutation

The most striking finding that emerged from the analysis of more than 1000 SARS-CoV-2 genomes was the preponderance of C->U transitions compared to other substitutions in the initial 4-5 months of its evolution. These accounted for 38%-42% of all changes in the four SARS-CoV-2 datasets. In seeking alternative, non-biological explanations for this observation, they cannot have arisen through misincorporation errors in the next generation sequence methods used to produce the dataset because the analysis in the current was restricted to consensus sequences. These are generally assembled from libraries that typically possess reasonable coverage and read depth; error frequencies of < 10^−4^ per site (14) would therefore improbably create a consensus change in a sequence library. There was furthermore, no comparable increase in G->A mutations (Fig. 2) and the sequence context in which sequencing errors occur (a 5’ or 3’ C or G; (14)) did not match the favoured context for mutation observed in our dataset (Fig. 6).

This asymmetric mutation furthermore cannot have arisen through a mutational effect of the coronavirus RNA dependent RNA polymerase during virus replication. By definition, a coronavirus RNA genome descends from any other through an equal number of copyings of the positive and negative strands – any tendency to misincorporate a U instead of a C would be reflected in a parallel number of G->A mutations where this error occurred on the minus strand (or vice versa). As demonstrated however, G->A mutations occurred at a much lower frequency than C->U mutations and were similar to those of A->G (Figs. 2A, 6).

The most cogent explanation for C->U hypermutation is the action of RNA editing processes within the infected cell. A well characterised antiviral pathway involves the interferon-inducible isoform of adenosine deaminase acting on RNA type 1 (ADAR1)(15). This edits A to inosine in regions of viral double-stranded RNA, which is subsequently copied as a G. Irrespective of its widely demonstrated antiviral role in a range of typically minus stranded RNA viruses, the mutations it creates do not match those observed in SARS-CoV-2 or other coronaviruses. Firstly, ADAR1 target dsRNA so editing effects tend to be symmetric with A->G substitutions being matched by T->C mutations. Secondly the direction of mutation is wrong. The focus of the analysis in the current study was on infrequent or unique polymorphisms where ancestral and mutant bases can be inferred. The excess of C->U transitions is the opposite of those induced by ADAR1.

A second, interferon-inducible pathway edits retroviral DNA during transcription and is strand specific; its typical antiviral activity is to mutate single stranded proviral DNA formed after first strand synthesis from genomic RNA (16-18). The deamination of C’s to T’s leads to the observed excess of G->A changes in the complementary positive stranded RNA virus genome (19). This editing function is performed by members of the apolipoprotein B mRNA-editing enzyme, catalytic polypeptide-like (APOBEC) family, many of which possess defined antiviral functions against retroviruses, hepatitis B viruses, small DNA viruses and intra-cellular mobile retroelements (reviewed in (20)). The APOBEC3 gene family members that are primarily involved in antiviral defence show evidence of extensive positive selection and expansion over the course of mammalian evolution, particularly in the primate lineage. Humans possess 7 active antiviral proteins (A3A, A3B, A3C, A3D, A3F, A3G and A3H) that contrasts with single A1 gene in rodents and a range of other mammals (21, 22).

While deamination of cytidines in single-stranded DNA sequences is a hallmark of APOBEC function, APOBECs show binding affinities for single stranded RNA templates that may mediate antiviral functions. A3B and A3F has been shown to block retrotransposition of a LINE-1 transposons mRNA through a non-deamination pathway (23), potentially through binding to single-stranded RNA. Direct editing of HIV-1 RNA by the rat A1 APOBEC and the accumulation of C->U hypermutation verified that RNA could also be used as a substrate for deamination (24). This suggested to the authors at that time that APOBEC-mediated RNA editing was a potential antiviral activity mechanism against RNA viruses as well as retroviruses.

Since then, evidence supporting this conjecture has been difficult to obtain; the virus inhibitory effect of APOBECs against enterovirus A71, measles, mumps and respiratory syncytial viruses were not shown to be associated with the development of virus mutations (25, 26). Similarly, A3C, A3F or A3H, but not A3A, A3D and A3G were shown to inhibit the replication of the human coronavirus, HCoV-NL63, but their expression did not lead to *de novo* C->U (or G->A) mutations on virus passaging (27). On the other hand, It has been demonstrated that A3A and A3G possess potent RNA editing capability on mRNA expressed in hypoxic macrophages (28), natural killer cells (29) and transfected A3G-overexpressing HEK 293T cells (30). These latter findings verify that APOBECs do possess RNA editing capabilities but do not provide any mechanistic context for the potential inhibition of RNA virus replication by this mechanism. Nevertheless the pronounced asymmetry in C->U transitions in SARS-CoV-2 and the preferential substitution of C’s flanked by U and A bases on both 5’ and 3’ sides (Fig. 6) that broadly matches what is known about the favoured contexts of A3A, A3F and A3H (31) provides strong circumstantial grounds for suspecting a role of one or more APOBEC proteins in coronavirus mutagenesis. Their exceptionally long genomes (≈30,000 bases), the exposure of genomic RNA in the cytoplasm before initiation of replication complex formation and the potentially lethal effects of just one mutations introduced into the genomic sequence makes APOBEC-mediated anti-coronaviral activity plausible in virological terms. It is perhaps because of the otherwise low mutation rate of coronaviruses and the extensive dataset of accurate, minimally divergent SARS-CoV-2 sequences assembled post-pandemic that has enabled this mutational signature to be so clearly observed.

### Evolutionary implications

The key finding in the study was the combined evidence for an APOBEC-like editing process driving initial sequences changes in SARS-CoV-2 and that the observed substitutions have not arisen through a typical pattern on random mutation and fixation that is assumed in evolutionary models. A specific problem for evolutionary reconstructions would be the existence of highly uneven substitution rates at different sites; APOBEC-mediated editing (and indeed the pattern of C-U transition in SARS-CoV-2 sequences) is strongly dependent on sequence context and, for at least two APOBECs, additionally influenced by their proximity to RNA secondary structure elements in the target sequence (28, 31). Sequence changes in SARS-CoV-2 and other coronavirus genomes may therefore be partially or largely restricted to number of mutational hotspots that may promote convergent changes between otherwise genetically unlinked strains. As demonstrated in Fig. 4, these can conflict with relationships reconstructed from phylogenetically informative sites. Furthermore, the substitution rate reconstructed for SARS-CoV-2 and potentially other coronaviruses may represent an uncomfortable amalgam of both the accumulation of neutral changes and forced changes induced by APOBEC-like editing processes that may obscure temporal reconstructions. A recent analysis of SARS-CoV-2 genomes illustrates these problems (https://doi.org/10.1101/2020.03.16.20034470); only a tiny fraction of variable sites (0.34%) were found to phylogenetically informative, while a high frequency of unresolved quartets demonstrates further the lack of phylogenetic signal in SARS-CoV-2 evolution reconstructions. The occurrence of multiple of driven change under host-induced selection is consistent with these cautionary observations.

The other important consequence of C->U hypermutation is that most of the amino acid sequence diversity observed in SARS-CoV-2 strains originates directly from forced mutations and therefore cannot be regarded in any way as adaptive for the virus (Fig. 5). An RNA editing mechanism of the type discussed above evidently places a huge mutational load on SARS-CoV-2 that may underpin the abnormally high dN/dS ratios recorded in SARS-CoV-2 and SARS-CoV sequence datasets (Fig. 1). It is likely that many or most amino acid changes are mildly deleterious and transient; repeated rounds of mutation at favoured editing sites followed by reversion may therefore contribute to the large numbers of scattered substitutions in SARS-CoV-2 sequences that conflict with their phylogeny.

Finally, it is intriguing to speculate on the long-term effects of the C->U/U->C asymmetry and the extent to which this may contribute to the previously described compositional abnormalities of coronaviruses (32, 33). As described above in connection with mutation frequencies, the compositional asymmetries cannot directly arise through viral RdRp mutational biases because any resulting base frequency differences would be symmetric (*ie*. G≈C, A≈U). Instead it appears that the observed imbalances in frequencies of complementary bases reflect the progressive depletion of C residues and accumulation of U’s by the APOBEC-like mutational process on the genomic (+) strand of coronaviruses. Culminating in the compositionally highly abnormal HKU1 sequences (32), this appears to have driven down the G+C content of coronaviruses as low as 32% while remarkably leaving G and A frequencies more or less unaltered (Fig. 7). Intriguingly, the bat-derived coronaviruses along with the recently zoonotically transferred viruses into humans show the least degree of compositional asymmetry.

The expansions in APOBEC gene numbers, extensive positive selection and the consequent variability in APOBEC nucleic acid targeting (20) may indeed create distinct selection pressures on coronaviruses in different hosts. The immediate appearance of C->U hypermutation in SARS-CoV-2 and SARS-CoV genomes in humans may therefore represent the initial effects of replication in a more hostile internal cellular environment than to be found in a better co-oadapted, virus-tolerised immune system of a bat (34). Zoonotic origins are suspected for other human coronaviruses but at more remote times (35); perhaps they have taken their mutational and adaptive journeys already.

## MATERIALS AND METHODS

### SARS-CoV-2 and other coronavirus datasets

The 1000 closest matched sequences to the prototype strain of SARS-CoV-2, NC_045512, were downloaded on the 24^th^ April, 2020. Sequences with large internal gaps, ambiguous bases and other markers of poor sequence quality were excluded, leaving a total of 865 sequences for analysis. This was divided into three data samples, corresponding to sequences 1-300, 301-600 and 601-884. A further dataset of 117 well curated SARS-CoV-2 sequences was downloaded from Konsiliarlabor für Coronaviren (https://civnb.info/sequences/) on the 13/04/2020 and represents a further independent sample set. A listing of further datasets of SARS-CoV, MERS-CoV and other human coronaviruses is provided in Table 1. All sequence alignments including accession numbers are provided as Supplementary Datasets.

### Sequence analysis

Calculation of pairwise distances and nucleotide composition, listing of sequence changes were performed using the SSE package version 1.4 (http://www.virus-evolution.org/Downloads/Software/) (36).

## ACKNOWLEDGEMENTS

The work was supported by a Wellcome Investigator Award Grant WT103767MA.

