## Supplementary Data Fig. S1 for "Rampant C->U hypermutation in the genomes of SARS-CoV-2 and other coronaviruses – causes and consequences for their short and long evolutionary trajectories"

FIGURE S1

FREQUENCIES OF TRANSITIONS IN THE 16 5’ AND 3’ BASE CONTEXTS

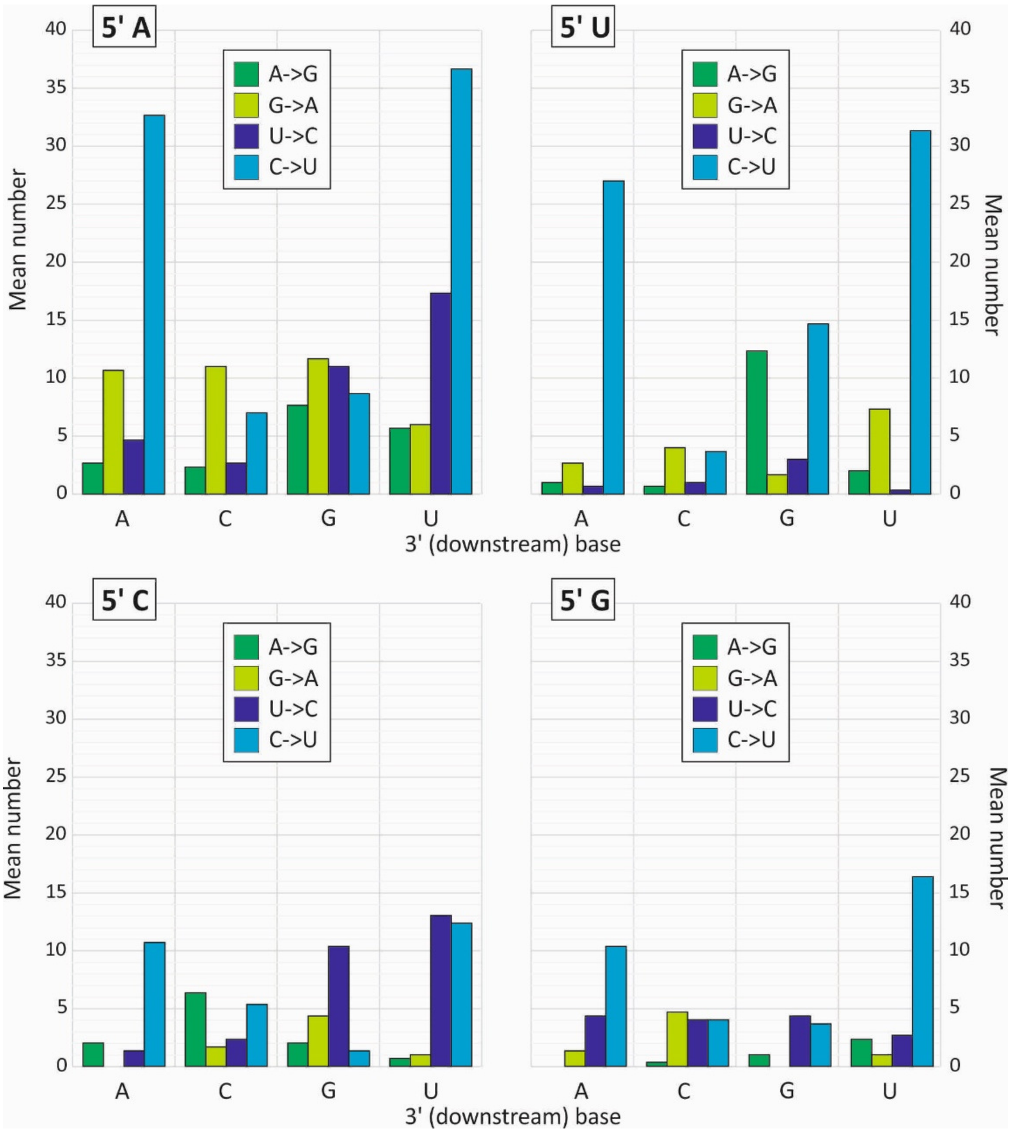


Effects of 5’ and 3’ bases on transition frequencies in SARS-CoV-2 full genome sequences.
